# The Influence of Neural Activity and Neural Cytoarchitecture on Cerebrovascular Arborization: A Computational Model

**DOI:** 10.1101/2022.03.14.484189

**Authors:** Bhadra S Kumar, Sarath C Menon, R G Sriya, V Srinivasa Chakravarthy

## Abstract

Normal functioning of the brain relies on a continual and efficient delivery of energy by a vast network of cerebral blood vessels. The bidirectional coupling between neurons and blood vessels consists of vasodilatory energy demand signals from neurons to blood vessels, and the retrograde flow of energy substrates from the vessels to neurons, which fuel neural firing, growth and other housekeeping activities in the neurons. Recent works indicate that, in addition to the functional coupling observed in the adult brain, the interdependence between the neural and vascular networks begins at the embryonic stage, and continues into subsequent developmental stages. The proposed Vascular Arborization Model (VAM) captures the effect of neural cytoarchitecture and neural activity on vascular arborization.

The VAM describes three important stages of vascular tree growth: (i) The prenatal growth phase, where the vascular arborization depends on the cytoarchitecture of neurons and non-neural cells, (ii) the post-natal growth phase during which the further arborization of the vasculature depends on neural activity in addition to neural cytoarchitecture, and (iii) the settling phase, where the fully grown vascular tree repositions its vascular branch points or nodes to ensure minimum path length and wire length. The vasculature growth depicted by VAM captures structural characteristics like vascular volume density, radii, mean distance to proximal neurons in the cortex. VAM-grown vasculature agrees with the experimental observation that the neural densities do not covary with the vascular density along the depth of the cortex but predicts a high correlation between neural areal density and microvascular density when compared over a global scale (across animals and regions).

To explore the influence of neural activity on vascular arborization, the VAM was used to grow the vasculature in neonatal rat whisker barrel cortex under two conditions: (i) Control, where the whiskers were intact and (ii) Lesioned, where one row of whiskers was cauterized. The model captures a significant reduction in vascular branch density in lesioned animals compared to control animals, concurring with experimental observation.

## Introduction

The neurons in the brain and the cerebrovascular network form a functionally integrated network [1]. Though the branching and patterning of neural and vascular arbours begin in the foetal stage itself [2], the two networks are capable of further growth, remodelling and plasticity postnatally also [3,4]. Electrical activity of the neurons in the brain is highly dependent upon timely and adequate blood supply from the cerebral vascular network; the cerebral blood flow, in turn, is dependent on the vasodilatory signals arising from the neural activity. Therefore, it is understood that there is a bidirectional functional connectivity between the neural and the cerebrovascular system. Hence, there is an anticipation, supported by experimental evidence [5–7], that even the development of both neural and cerebrovascular systems should be interdependent [8–10].

Development of the cerebrovascular network is driven by two processes, namely, vasculogenesis and angiogenesis [11]. Vasculogenesis is the formation of new vessels from the vascular precursor cells. In contrast, angiogenesis is the formation of new vessels from the existing vessels by a process of branching and arborization, which forms an intricate vascular network. In the embryonic stage, shortly after the closure of the neural tube, between the embryonic day 8.5 (E8.5) and E10, a primitive vascular network is formed around the neural tube. This primitive network of vessels, called the peri-neural vascular plexus (PNVP), develops in response to the vascular endothelial growth factor A (VEGF-A) released by the neural tube [7,12,13]. The proangiogenic nature of VEGF-A activates the vasculogenesis of the neural tube. A variety of cellular and molecular pathways are involved in neurally-guided cerebral angiogenesis [7,8,10]. In the post-natal stage too, angiogenesis continues to take place and is known to be influenced by the VEGF of neuronal origin [14]. Thus, it can be stated with reasonable confidence that the vascular arborization depends on the neural cytoarchitecture and the preliminary arborization of vessels can be modelled simply using the neural cytoarchitecture data.

The neural architecture and organization in many brain areas indicate the tendency of minimizing wiring costs in the development of neural circuitry [15–18]. Cuntz and colleagues [19,20] have shown that a variety of dentritic arborization topologies can be accounted for by the minimum wiring cost principle, a principle that is implemented in the form of a simulation system known as the TREES toolbox [19,20]. The work of [19] presents a simple algorithm which, by optimizing path length and wirelength, can generate structures strikingly similar to a variety of biologically observed neurons. We propose that the vascular arborization also might be aiming for optimally spanning the tissue volume while ensuring minimal path length and wire length. Given the neural distribution and the vascular growth principle, we show how we can simulate the tree growth architecture that closely matches the experimentally observed vascular arborization characteristics.

Several computational approaches were proposed to model vasculature [21–25]. But to the best of our knowledge, there are no computational models that describe the growth and spanning of cerebro-vasculature based on the neural distribution and neural activity. In response to this lacuna, we propose a computational model known as the Vascular Arborization Model (VAM) to understand the effect of activity and density distribution of the neurons on the development of the vascular arborization

In the first half of the work, we simulate the vascular tree growth in a small volume (0.1mm X 0.1mm X 0.8mm) of the cortex. The vascular network originates from a parent root node and grows to span the tissue volume based on the cytoarchitecture of the neural and non-neural cells of the cortex. There are two stages of vascular arborization: (i) Growth phase and (ii) Settling phase. In the growth phase, each terminal vessel grows closer to the neurons in its domain such that the mean squared distance between the vascular terminal node and the neurons in its perfusion domain is minimized. When the mean squared distance reaches a minimum, the terminal vessel splits into two smaller vessels. The former perfusion domain is also split into two sub-domains, while the two child terminal vessels now proceed to perfuse the two newly formed subdomains. This process occurring iteratively results in vascular arborization. During the settling phase, the fully grown vascular network is rearranged such that the vascular tree ensures optimal path length and wire length.

In recent studies, it was observed that in addition to the location of neural structures, the neural activity also could influence the vascular patterning [5,6,26]. Taking this into consideration, in the second half of the work, we modified the VAM model to allow the arborization of vessels depend on neural activity as well. We used a Self-Organizing Map (SOM) model [27] to simulate the neural map responses and use it to map the whisker inputs. This neural response was used as the activity in layer 4 (L4) of the whisker barrel cortex. The growth phase was made dependent on neural activity also such that the vessels would not sprout until a minimum neural activity is present in its perfusion domain. The settling phase also was made to depend on the neural activity. We observed the branch point density and vascular volume density in a control model and a lesioned model. The model simulations were close to the experimental observations [6].

## Methods

The VAM was implemented for a small 3D volume of the somatosensory cortex of the murine brain. The neuronal density distribution of the murine cortex observed experimentally [28] was used to distribute the neurons along the depth of the cortex in the VAM model. The vasculature was initialized as a set of randomly distributed root nodes at the surface of the cortex within the considered volume.

In the phase 1 of the simulation study, we simulate the vascular arborization based on cytoarchitecture of neurons and non-neuron cells, and verified the results obtained using the vascular volume density observed experimentally. Here we considered a smaller section of the somatosensory cortex of dimension 100μm X 100μm X 800μm which has one single root node.

In the second part of the model, we considered a larger area of the somatosensory cortex 600μm X 600μm X 700μm with 44 root nodes, and incorporated the neural activity in L4 to make the arborization depend on the activity in addition to cytoarchitecture.

### Simulation of neural distribution

The neural cell number density along the depth of the cortex was obtained from the fig.10 (a-d) of Tsai et al 2009 [28]. In phase 1, a volume of dimension 100μm X 100μm X 800μm was considered, whereas in phase 2, a volume of dimension 600μm X 600μm X 700μm was considered. The volume was segmented into smaller layers, each of thickness 50μm (dimensions of individual layer in phase 1 are 100μm X 100μm X 50μm, and in phase 2 it is 600μm X 600μm X 50μm) stacked one on top of another along the radial direction of the cortex. The number of neurons in each layer was estimated based on the experimental values of cell number density of neurons at each 50μm depth. Once the number of neurons in each layer was estimated, they were distributed randomly in the layer.

#### I. The vascular tree arborization algorithm based on cytoarchitecture

The proposed the vascular tree arborization algorithm consists of two phases:

Phase I: Growth phase,
Phase II: Settling phase.

In Phase I, the tree growth is based on the cytoarchitecture of the neural and non-neural cells (hereafter called “cortical cells”) of the cortex. The vascular tree starts as a single child node starting from a root node. During the growth phase, a single vascular node, emerging from the root node, grows incrementally and splits into branches at appropriate locations ensuring minimum distance between the terminal vessels and the tissues to be perfused. During the settling phase, all the vascular nodes are repositioned such that the tree has an optimal path length and wire length. By pathlength, we define the total distance along the path traced from a leaf node to the root node (in fig. 2.a, the blue arrows from leaf node to root node trace one path). The wirelength simply denotes the total length of all the vascular segments in the vascular tree (in fig. 2.a, each green arrow represents a wire).

##### Phase I: The growth phase

The growth phase describes the process where the rudimentary vascular tree, consisting of a single vascular node, originating from the root node at the cortical surface, grows out, branches and perfuses the tissue volume. The tree which starts its growth from the root node (fig.1.a), incrementally grows and spans the space of the tissues such that the average distance between the energy demanding cells (neurons and non-neural cells) and the vascular leaf nodes is minimum.

**Figure 1:**
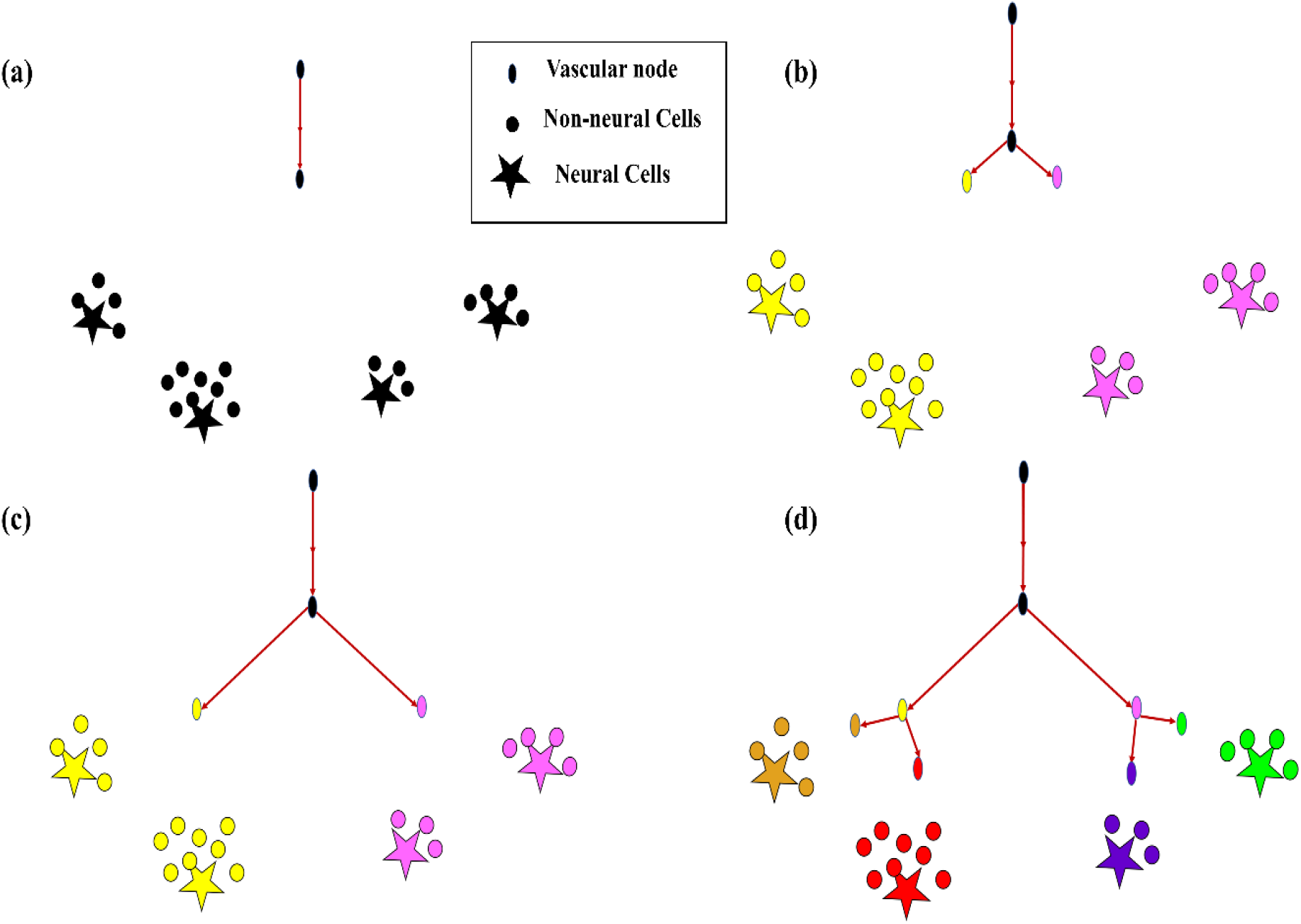
The growth of vascular tree from a root node. (a) All cortical cells (neurons and non-neural cells) fed by a single vascular leaf node (black); (b) The earlier leaf node (black) splits into two leaf nodes (yellow and violet), splitting its perfusion domain also into two regions of cortical cells assigned to the yellow and violet vascular leaf nodes; (c) The yellow and violet vascular leaf nodes move towards the centroids of their respective perfusion domains; (d) Once the equilibrium point is reached, the yellow node splits again into two leaf nodes (gold and red) and the violet node splits into two leaf nodes (blue and green), splitting their perfusion domains also into four regions corresponding to the four terminal leaf nodes (gold, red, blue and green).

The total number of cortical cells (neurons and non-neurons) is fixed to be *N_T_* throughout the simulation, and the total number of nodes in the vascular tree are *M_T_*(*N_T_* > *M_T_*). At any given instant during the tree growth, let the set of leaf nodes (terminal vascular nodes) be denoted by LN. Each cortical cell is assigned to the closest node (minimum Euclidean distance) among the *m* leaf nodes. Given that the coordinate of the j^th^ leaf node is given by ***Q_j_*** and the coordinate of the i^th^ cortical cell is given by ***P_i_***, the sum of squared distance between the i^th^ cortical cell and j^th^ leaf node which is the leaf node closest to the i^th^ cortical cell, denoted by j(i), at any instant is given by,

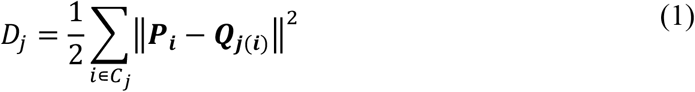

where *C_j_* is the set of cortical cells assigned to the j^th^ leaf node.

The total distance between the vascular leaf nodes and the cortical cells is given by,

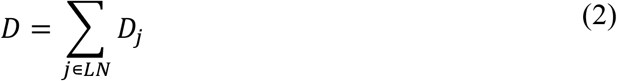

The jth leaf node is moved closer to the cortical cells such that *D* is minimized

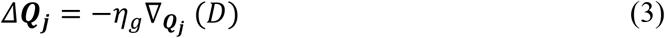

where *η_g_* is the learning rate.

Given that ***Q_j_*** is represented by a 3-dimensional vector 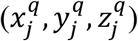

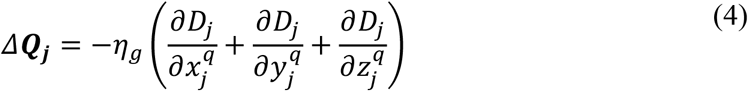

This gradient is calculated for every *j* = 1: *m*. The jth leaf node once reaches a point such that *Δ**Q_j_*** ≈ 0, then the jth node splits giving rise to k child nodes which would be the new leaf nodes. Now the cortical cells are reassigned to the newly formed leaf nodes (fig. 1b and 1d) based on the nearest distance criterion. The new leaf nodes continue to grow like before. The tree growth terminates when the number of vascular terminal leaf nodes count up to a predefined fraction of the total neurons. That is, the branching stops when *m* = *n* * *α*, where *α* ≤ 1. The level (*l*) of a vessel is identified by the number of branchings that occurred from the root node.

##### Phase II: The settling phase

Once the tree spans the space such that there are m leaf nodes, the settling phase begins. Here, a grown tree rearranges itself so as to optimize the cost of path length and wire length. To optimize the path and wire lengths, we try to minimize the sum of squared pathlengths, sum of squared wirelengths and sum of squared distances from leaf nodes to tissues. The cost function to be optimized consists of three components, (i) Total squared path length (TSPL), (ii) Total squared wire length (TSWL), (iii) Total squared tissue-vascular distance (TSTD).

For a given vascular tree, let there be L number of levels. Let the level of hierarchy of any node j in the vascular tree be denoted by *l_j_*. The root node is identified as the top of the hierarchy with level number, *l_j_* = 1. Let the level number of the vascular terminal leaf nodes be *l_j_* = *L*.

1. Total squared path length (TSPL) minimization: The TSPL of the vascular tree is calculated as,

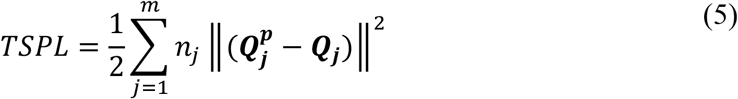

where *m* is the total number of nodes in vascular tree, 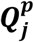 is the position of the parent node j, ***Q_j_*** is the position of the j^th^ node, *n_j_* is the number of leaf nodes downstream the node j (fig 2.b). The ***Q_j_*** are updated by performing gradient descent on TSPL as follows,

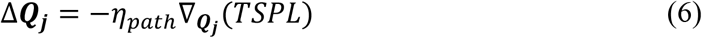
2. Total Squared Wirelength (TSWL) minimization: The TSWL of the vascular tree is calculated as

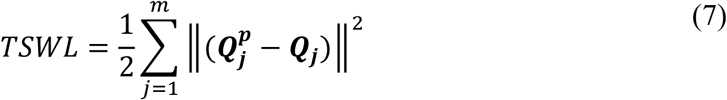

where *m* is the total number of nodes in vascular tree, 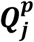 is the position of the parent node j, ***Q_j_*** is the position of the j^th^ node. The ***Q_j_*** are updated by performing gradient descent on TSWL as follows,

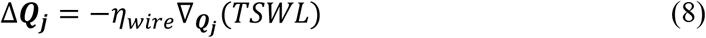
3. Total squared tissue-vascular distance (TSTD) minimization: TSTD estimates the total squared distance between the vascular nodes and the cortical cells assigned to them. The distance between a vascular leaf node j (j ∈ LN), and the cortical cells (***P_i_***, *i* ∈ ***C_j_***), where ***C_j_*** denotes the set of cortical cells assigned to the vascular leaf node j, is given by

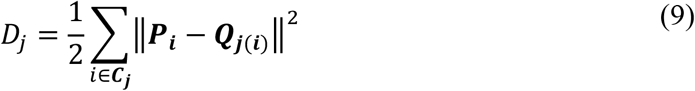 The TSTD is calculated as

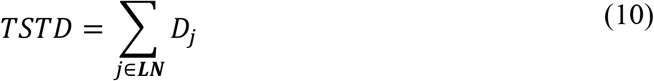

where **LN** denotes the set of leaf nodes. Note that the expressions on the right-hand side in eqns. (9,10) are the same as those in eqns. (1,2) respectively. The ***Q_j_’***s are updated by performing gradient descent on TSTD as follows,

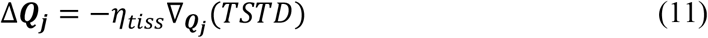 Jointly, eqns. (6, 8, 11), govern the settling process, which continues until the vascular tree converges. Convergence criterion is given by, *ξ* → 0, where *ξ* is defined as,

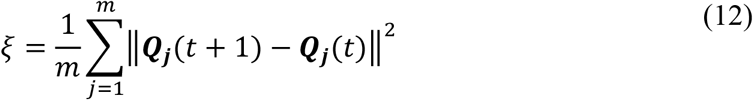

**Figure 2:**
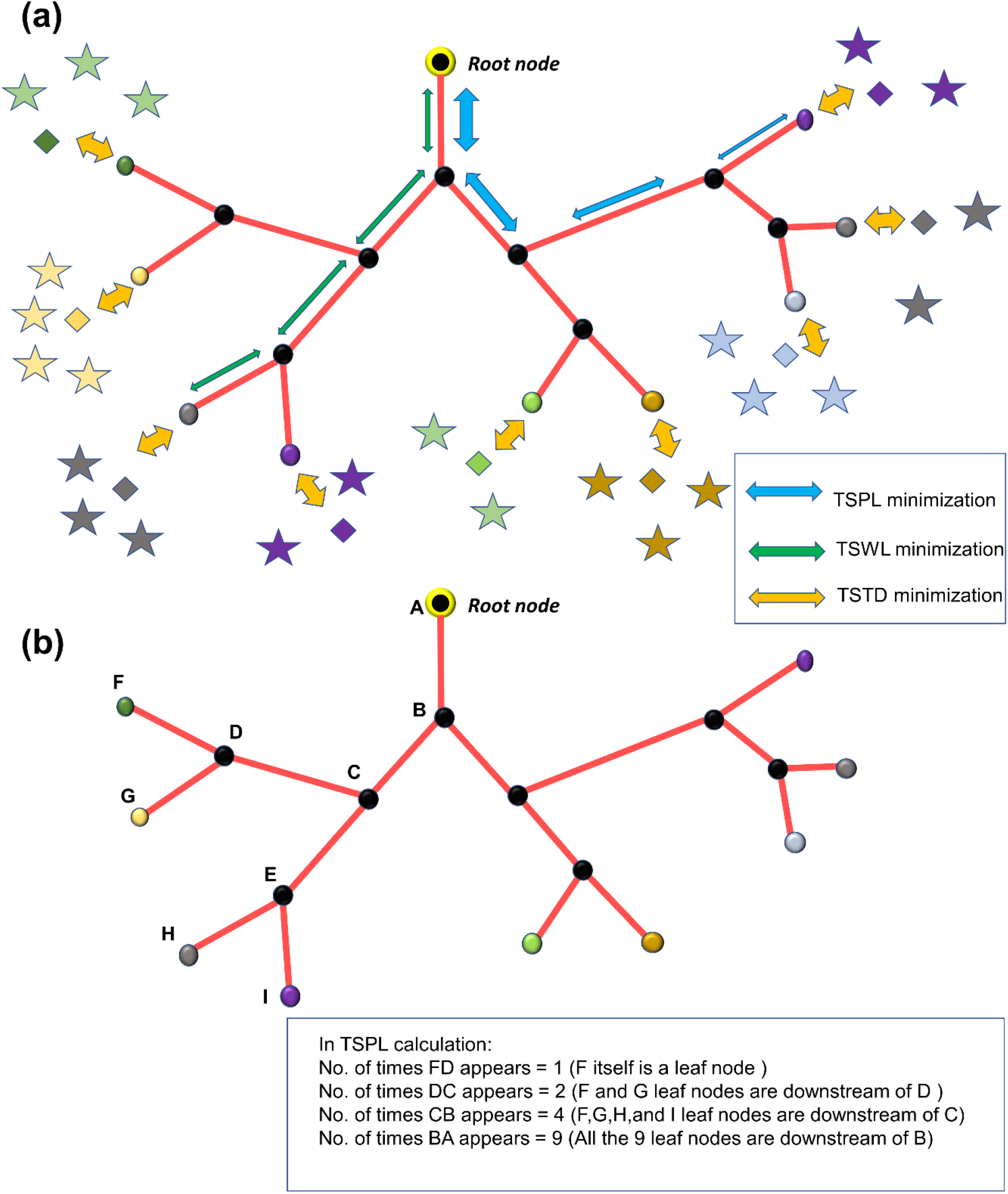
(a) The minimization of TSPL, TSWL, TSTD shown in the vascular tree; (b) estimation of *n_j_*, the number of leaf nodes downstream of the vascular node, j.

##### Estimation of the length of vessels

The length of a vessel formed by connecting a parent node x to a child node y, is defined as the Euclidean distance between the parent node (x) and the child node (y).

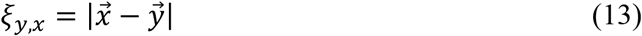

Since we estimate the volume density in each layer, the length contribution of each vessel in each layer is identified by measuring its length inside the specified layer.

##### Estimation of the vessel radius

The radius of each vessel is defined based on its level of hierarchy in the vascular tree. The radius at each level is iteratively fit in such a way so as to obtain the volume density distribution as close as possible to experimentally observed values. Two steps of iteration are followed. First, the microvessel radii are approximated to fit the microvascular volume density observed experimentally. Then, after fixing the microvascular radii, the radii of the higher levels are estimated such that the total vascular volume density is closest to the experimental observation. All radii below 3μm were considered to be microvessels. The level at which the vessels become microvessels was considered as a free parameter.

The radius of the vessels at each level of hierarchy in the vascular tree was arranged as an array (R) of dimension Lx1 (L is the number of levels) and was initialized as *R* = *R*_0_. The vascular volume density from the model 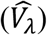 is estimated based on the value of R. The desired output (*V_λ_*) is the value of the vascular volume density at each layer *λ*, as obtained from the experiment [28]. The cost function (C) is defined by the mean squared error between *V_λ_* and 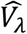.

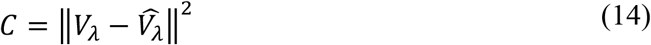

The radius of a randomly chosen level was increased by a small value ΔR such that C decreased continuously. Once an update is made to radii of all levels, the radii were adjusted to make sure that they follow a descending order of magnitude from root node level to leaf node level. The iterations were carried out until the magnitude of C was less than 10^−9^.

The approximation was also carried out using linear regression. The cost function was calculated for each layer (*λ*). In each layer the vessels were identified based on their level of hierarchy in the vascular tree, and a radius was assigned to each of them 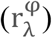 accordingly. The length contribution of each vessel to that layer was also calculated 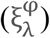 using trigonometric principles (Please see supplementary material section S1 for detailed explanation). The cost function per layer is defined by

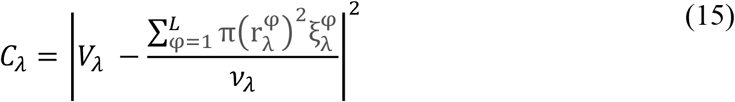

where *v_λ_* = Length X Width X Height of the volume of an individual layer. Note that in eqn. 15, we use the formula for the volume of a cylinder πr^2^ 1, where r is the radius and l the length of the cylinder.

The gradient of the radii of vessels at each level of hierarchy in the vascular tree was calculated such as to minimize the cost function (*C_λ_*)

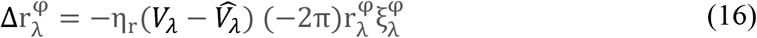

The iteration was looped over the total number of layers and the entire iteration was repeated until C was less than 10^−9^.

But since we consider stacked layers instead of sliding layers with overlap, both the above methods were only able to yield a crude approximation.

##### Estimation of volume density in each layer

Once the optimal values of radii for each layer were obtained, the vascular volume density 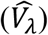 was calculated for each layer using the equation below.

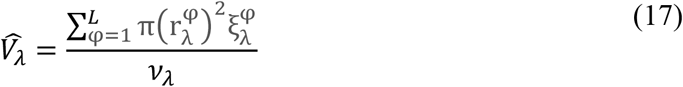

The microvascular volume density 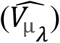 was calculated by considering the microvessels only (vessels whose radii is less than or equal to 3*μm*).

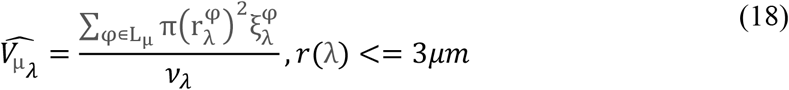

where L_μ_ indicates the level of hierarchy in the vascular tree beyond which the vessel radii are less than or equal to 3*μm*.

##### Estimation of mean distance between vessels and neurons

The distance between a neuron at coordinate ***P_i_*** and its closest vessel at coordinate ***Q***_***j***(***i***)_ was estimated for all neurons in *λ*’th layer. The mean squared distance (*D_λ_*) in that layer was defined as the average of squared Euclidean distance between all neurons in that layer and their respective closest vascular leaf node. Let the set *S_λ_* denote the indices of the neurons which belong to layer *λ*.

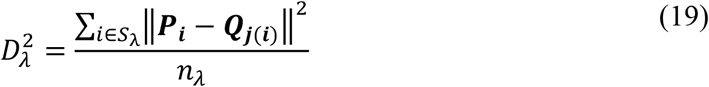

#### II. Vascular Arborization Based on Neural activity and Neural Cytoarchitecture

In the first part of the VAM simulation study, we simulated a small volume in the somatosensory cortex (100μm X 100μm X 800μm). The model was able to grow a vascular tree true to the experimental observations. Hence, we expand the model to a larger volume spanning a volume of (600μm X 600μm X 700μm). The number of root nodes in this larger volume was estimated to be ~44 as per the experimental observations by Wu et al [29]. The root nodes were fixed randomly from a gaussian distribution. The VAM model was modified so as to incorporate the influence of neural activity in addition to neural and non-neural cytoarchitecture on the vascular tree growth. The modified VAM model is called the Activity-based Vascular Arborization Model (A-VAM)

Two types of metrics are used to compare the A-VAM simulations with the experiments: (a) Branch Point Density (BPD), and (b) Vascular Length Density (VLD). The BPD is defined as the density of the vascular nodes in a given volume. Given that in a tissue volume (*ψ*) with length a, width b, and height c, the total number of vascular nodes counts to *m_ψ_*

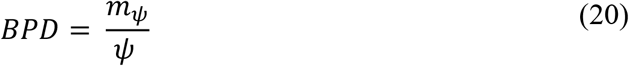

where *ψ* = *abc*, is the total volume of the tissue under consideration.

The Vascular Length Density (VLD) is defined as the ratio of the cumulative length (*α_ψ_*) of all the vessels in the volume to the volume of the tissue.

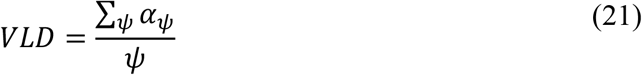

In A-VAM, there are three major stages for the tree growth (i) Prenatal growth phase, (ii) Post-natal growth phase, and (iii) Settling Phase.

##### (i) Prenatal Growth Phase

This phase simulates the vascular arborization which occurs during foetal development until the birth. Here we assume that there is minimal to nil neural activity and hence the vascular arborization completely depends on the neural cytoarchitecture. We run the prenatal growth phase to ensure that the BPD and VLD are close to experimentally observed values at birth.

##### (ii) Post-natal Growth Phase

During post-natal phase, we assume that the vascular arborization depends on neural activity in addition to neural cytoarchitecture. The neural network is simulated using a Self-Organizing Map (SOM) which results in a topographic mapping of input [27]. The SOM model is trained using input stimuli curated to imitate the whisker simulations. Each whisker simulation is represented as a two-dimensional Gaussian distribution centered around the pixel representing the whisker under simulation. The afferent weight connections of the winning neuron and its neighbours are moved closer to the input vector scaled by a Gaussian function which peaks at the winning neuron (please see supplementary material section S2 for details of SOM training).

The neural activity from whisker simulation is assumed to have the highest activity in layer 4 (L4) of the whisker barrel cortex ([29]). Hence in the model, we introduce the neural activity only in L4 (the region between depth 0.3 mm to 0.5 mm from the surface). The SOM layer is stacked up in this volume as shown in fig. 3. The neural units in the SOM are distributed uniformly in each layer as shown in the right panel in fig. 3 (white circles). The activity (*δ_i_*) of each i^th^ cortical neuron (red star in figure 3, right panel) depends on the activity of the nearest SOM neuron (*Z_i_*) and its distance (*ϑ*) from the SOM neuron as follows.

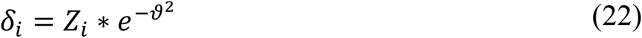

**Figure 3:**
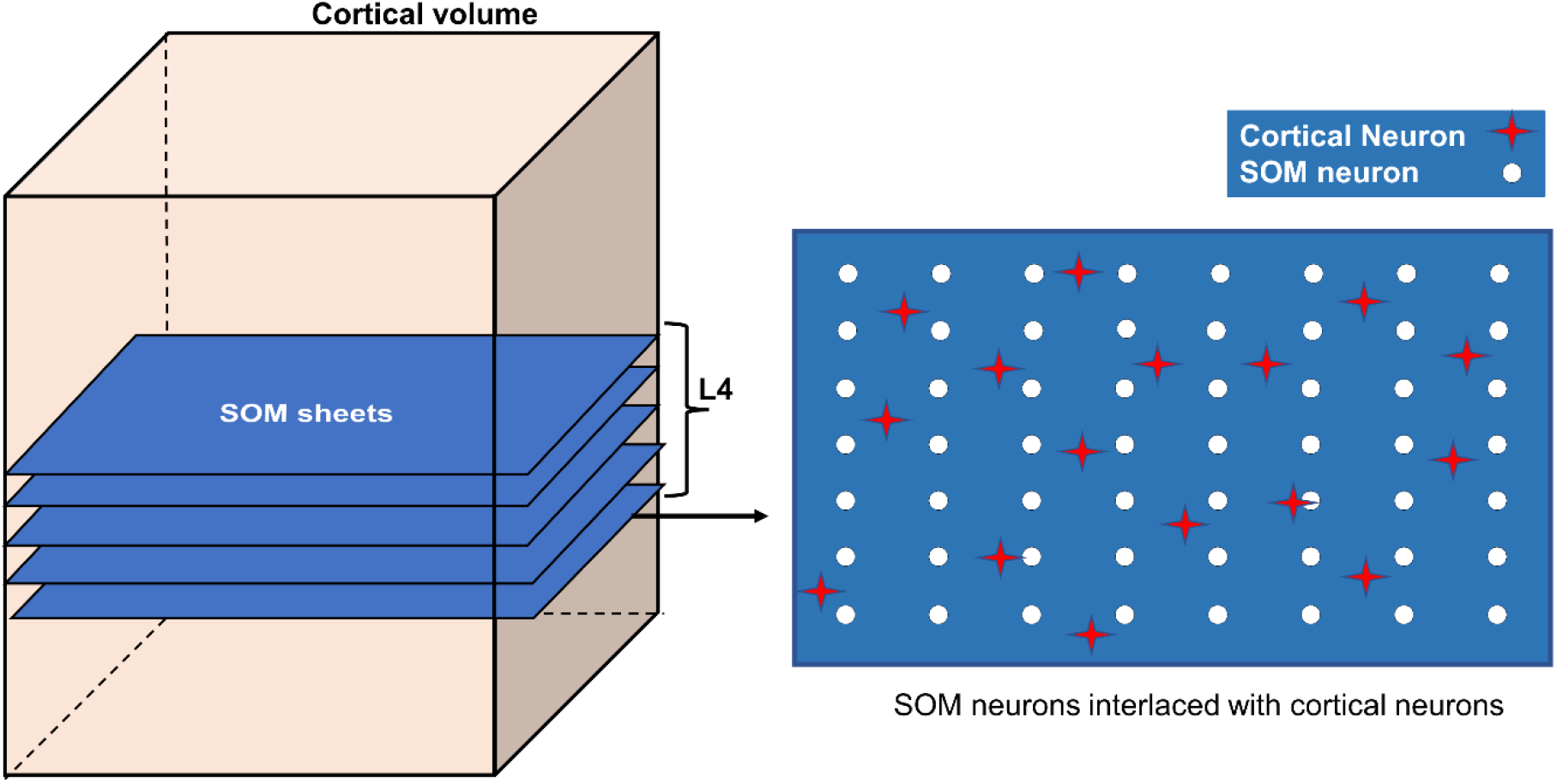
Incorporating SOM in VAM to form A-VAM

The vascular tree growth equation was modified such that the neural activity (*δ_i_*) also influences the learning rate of the growth and arborization of the vessels.

While incorporating the effect of neural activity in the growth phase, the learning rate, *η_g_*, varies for each neural tissue based on its activity. The learning rate (*η_g_i__*) at which each i^th^ neuron influences the calculation of gradient Δ***Q_j_*** of the j^th^ vascular node is given by

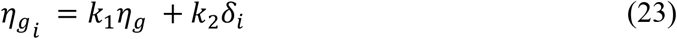

where *k*_1_ and *k*_2_ are constants and i∈Cj.

An additional constraint was also introduced to make the stopping criteria also depend on the neural demand. In order for a vascular node to split and give rise to child nodes, the perfusion domain of the vascular node should have a minimum neural activity (*δ_min_*) in addition to the requirement of Δ***Q_j_*** ≈ 0. If a node j has a very small gradient (Δ***Q_j_*** ≈ 0) but does not have the minimum neural activity in its perfusion domain, then that node is declared as a leaf node. The vascular network grows until all the nodes are labelled as terminal nodes.

##### (iii) The Settling Phase

The settling phase starts once the tree is grown completely. Similar to the settling phase in VAM, in A-VAM also, fully grown trees undergo settling under the influence of neural activity in addition to tissue cytoarchitecture. The learning rate (*η_tiss_*), used while calculating the gradient Δ***Q_j_*** using the TSTD, would depend on the neural activity as follows:

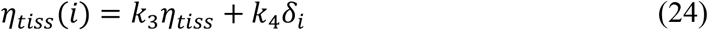

where i∈Cj.

| Table of parameters and constants |  |
| --- | --- |
| $\eta_g$ | $5*10^{-5}$ |
| $\delta_{min}$ | 0.07 |
| $\eta_{path}(prenatal)$ | $1.7*10^{-4}$ |
| $\eta_{wire}(prenatal)$ | $1.15*10^{-4}$ |
| $\eta_{tiss}(prenatal)$ | $2.5*10^{-4}$ |
| $\eta_{path}(postnatal)$ | $1.8*10^{-4}$ |
| $\eta_{wire}(postnatal)$ | $1.5*10^{-4}$ |
| $\eta_{tiss}(postnatal)$ | $2*10^{-4}$ |
| $k_1$ | 0.1 |
| $k_2$ | $1*10^{-4}$ |
| $k_3$ | 1 |
| $k_4$ | $9*10^{-5}$ |

## Results

### Distribution of neurons

The neural distribution was simulated for four different regions considered in the experiment by Tsai et al [28]. The four regions were: (a) Agranular cortex which was measured 1mm anteroposterior and 1mm mediolateral of the Bregma, (b) Partially granular cortex measured - 1 mm anteroposterior and 1 mm mediolateral of the Bregma, (c) Partially granular cortex measured 2 mm anteroposterior and 1 mm mediolateral of the Bregma, and (d) Granular cortex measured 1 mm anteroposterior and 4 mm mediolateral of the Bregma. The model assumed 16 layers of 50 μm height stacked one on top of another and sampled the neural cell density at each 50μm depth to reconstruct a similar neural distribution. The reconstructed neural cell number density is shown by the red plots in figures 4a to 4d. The experimental values were obtained from the fig.10 of [28].

**Figure 4:**
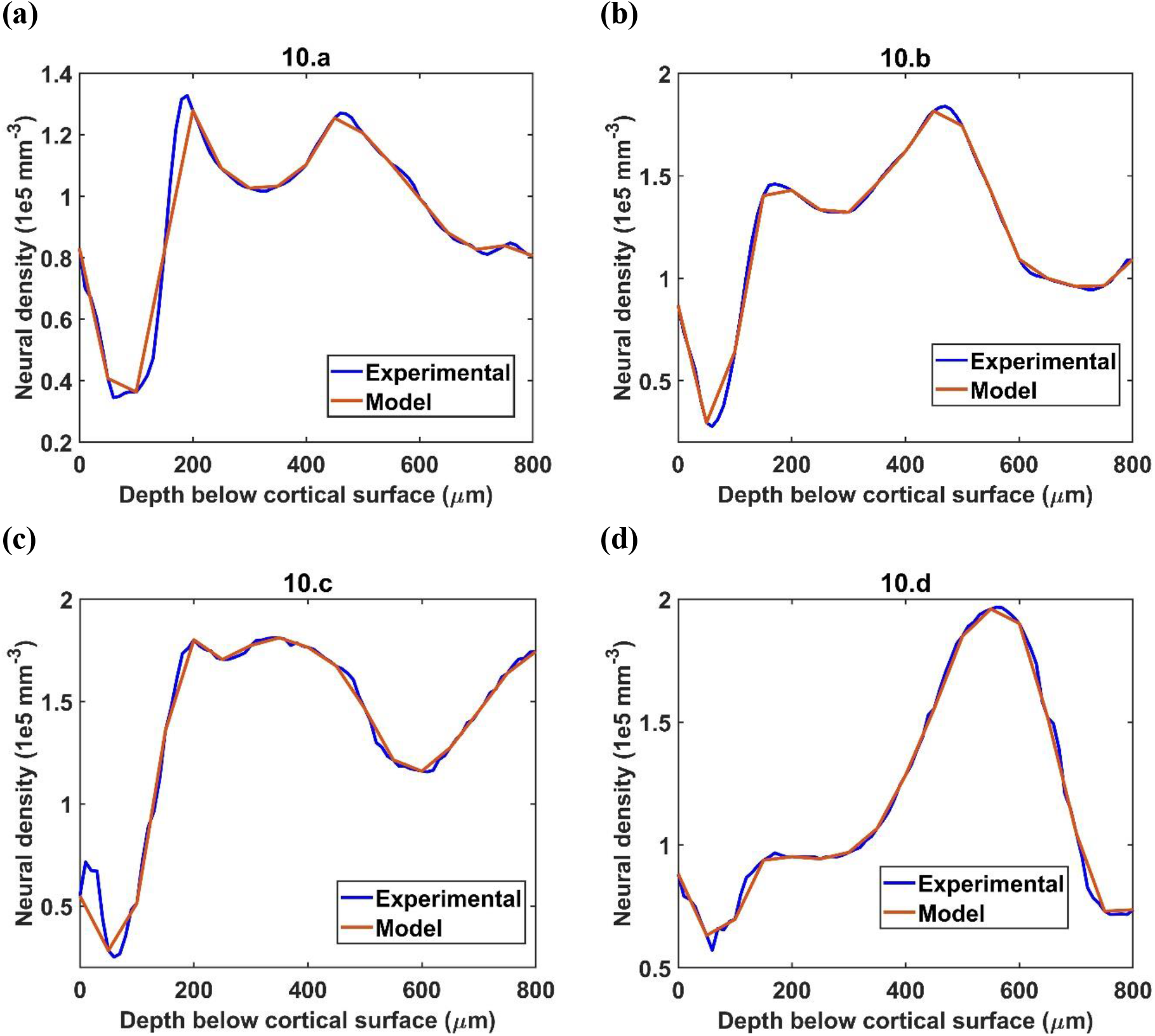
The simulation of neural distribution along the depth of the cortex in 4 different regions. The title shows the reference figure number in Tsai et al 2009 which was reconstructed by the model. The red plot shows the reconstructed neural cell distribution from the model.

### Vascular tree growth

An example of neural distribution inside a slab of dimension 0.1 mm X 0.1 mm X 0.8 mm is shown in fig. 5.a. The vascular node was initialized at the centre of the cortical surface as shown by the larger red sphere in fig. 5b. Fig. 5c shows the tree after the settling phase. Even though the difference in tree structure is not evident by a comparison of fig. 5b and fig. 5c, settling phase brings in fine adjustments in the tree which is evident clearly in the comparison of vascular volume density comparison.

**Figure 5:**
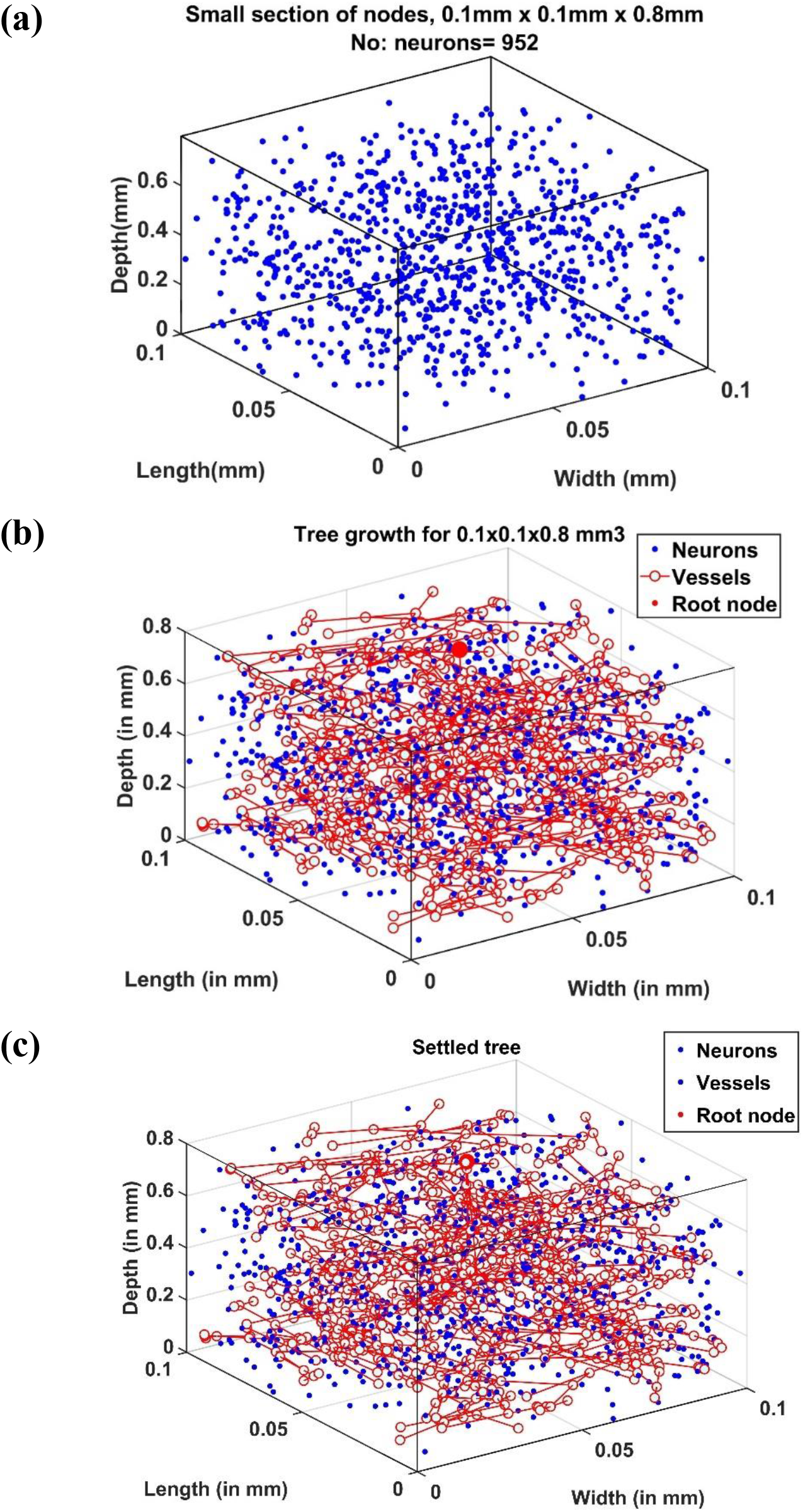
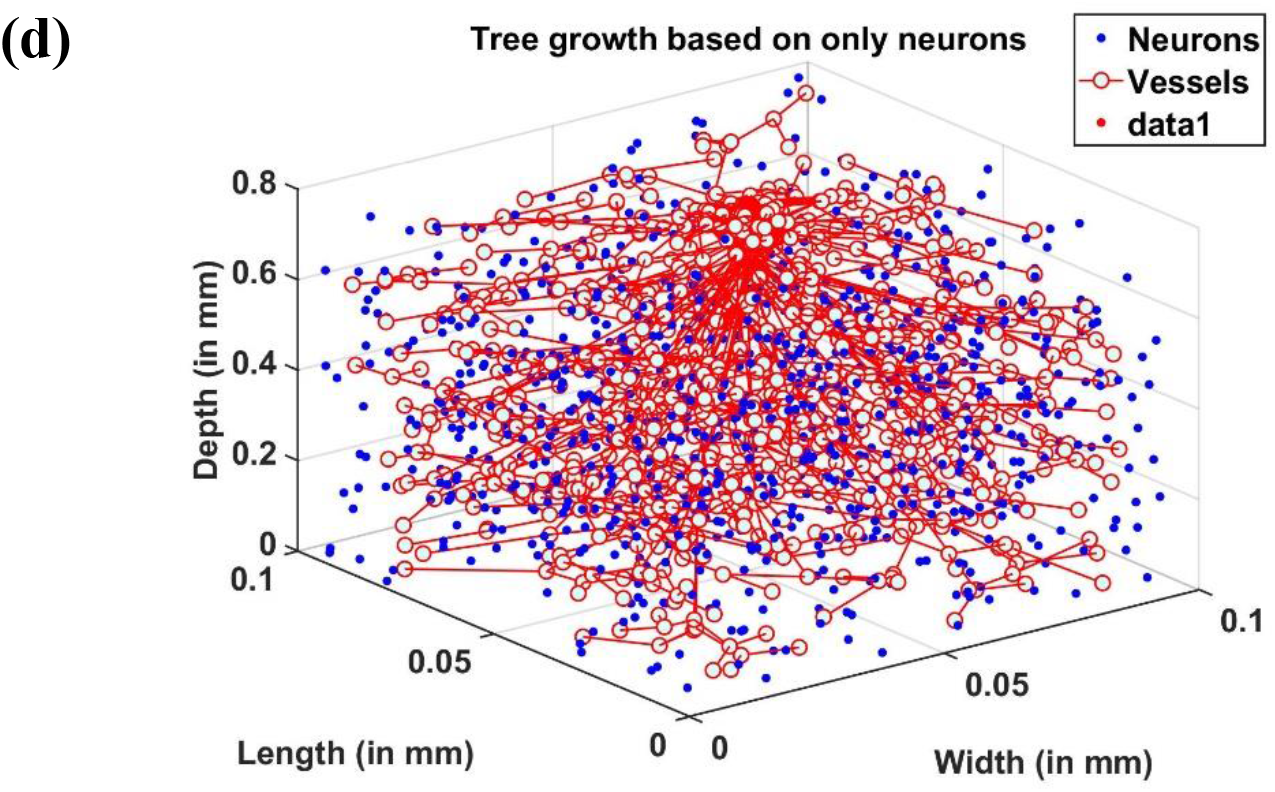
(a) Neural distribution in the slab of tissue under observation, (b) Tree formed after the growth phase (grown under the influence of all cortical cells), (c) Tree formed after the settling phase (grown under the influence of all cortical cells), (d) Tree grown and settled under the influence of only neurons.

The tree growth phase was tried using two conditions – the first one by assuming that the tree growth would be influenced by cytoarchitecture of neurons only. The tree grown based on this assumption is shown in fig. 5d. The alternate assumption was that not only neurons but non-neural cells also influence the tree growth. The grown tree in figs. 5b,c was based on this assumption. In both fig. 5b,c and fig. 5d, the number of vascular leaf nodes were fixed to be equal to 70% of the total number of non-neurons. This was to ensure that in both the assumptions, the number of leaf nodes are equal.

### Radius of the vessel at each level in vascular tree hierarchy

The radius of the vessel at each level was iteratively calculated to give the best approximation of the experimentally observed vascular volume density of the vessels (fig. 6). The range of radii are in agreement with the range of arteriolar diameter observed experimentally in rats and mice [30,31]. The maximum diameter of the blood vessel was 30μm. The radius decreases at each level of the bifurcation until the vessel is declared a terminal leaf node which is analogous to the capillaries whose diameter is close to 4μm (radius ~2μm) agreeing with experimental observations [11].

**Figure 6:**
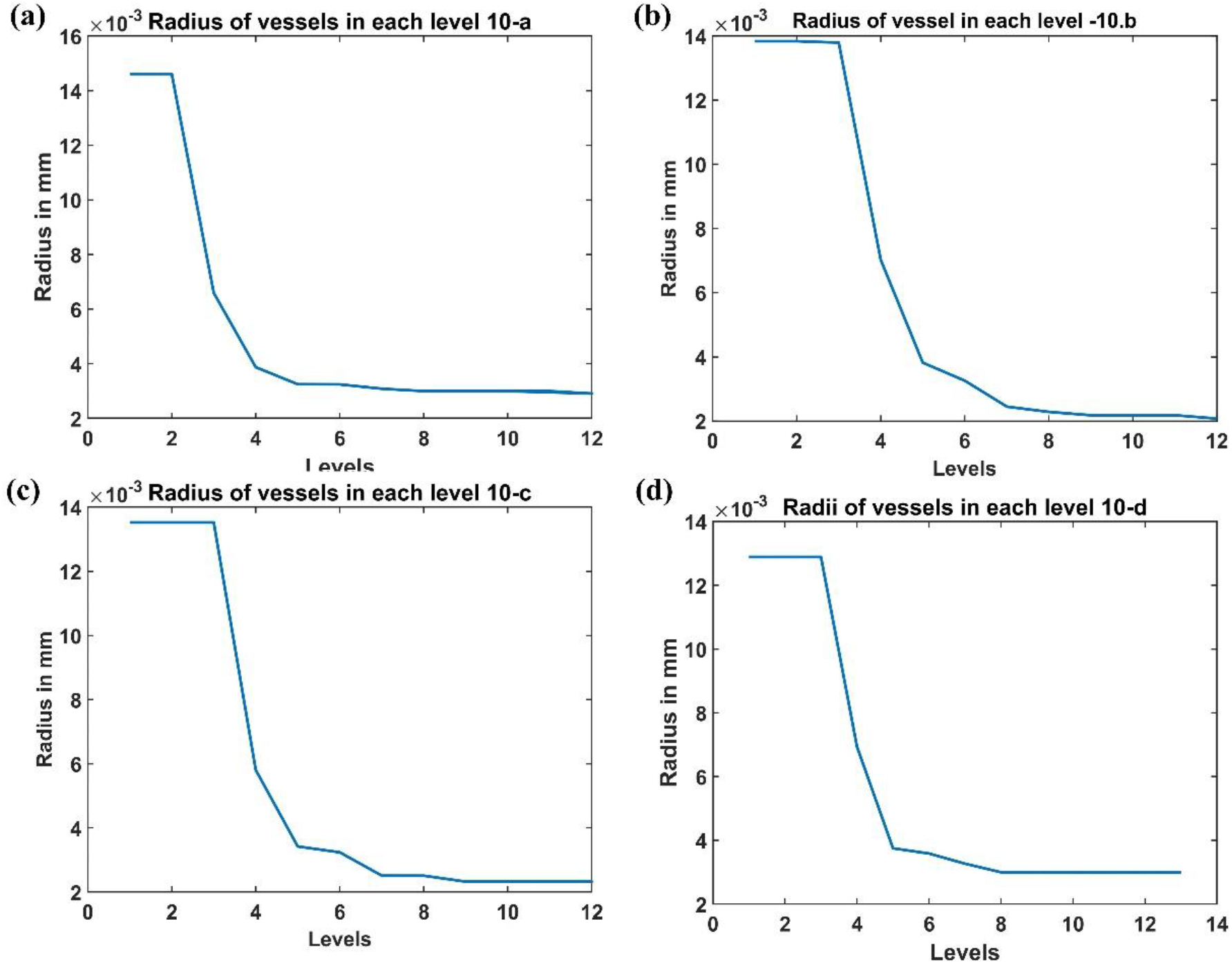
The radii at each level estimated from the A-VAM study for the following 4 regions: (a) Agranular cortex which was measured 1mm anteroposterior and 1mm mediolateral of the Bregma, (b) Partially granular cortex measured −1 mm anteroposterior and 1 mm mediolateral of the Bregma, (c) Partially granular cortex measured 2 mm anteroposterior and 1 mm mediolateral of the Bregma (d) Granular cortex measured 1 mm anteroposterior and 4 mm mediolateral of the Bregma

### Vascular volume density

The vascular volume density was calculated at each layer of dimension 0.1mm X 0.1mm X 0.05mm. The vessels in each layer were identified and the volume contribution was calculated based on its length in the layer and the level of the vessel. The figs. 7a-d compare the reconstructed vascular volume density in the four regions of the cortex. The vasculature was grown using two methods: (i) only neurons influence the vascular growth, and (ii) both the neurons and the non-neurons influence the vascular growth. As can be observed from the plot, the green dashed line shows the vascular volume density estimated from the settled tree grown using all the cells, the neurons and the non-neural cells. The blue dashed line on the other hand is the vascular volume density obtained from the tree grown using only neurons. We can observe that the estimation from the tree grown using all the cells is a better approximation than from the model grown based on only the neurons. This suggests that the cytoarchitecture of both neurons and non-neural cells plays an important role in deciding the vascular architecture.

**Figure 7:**
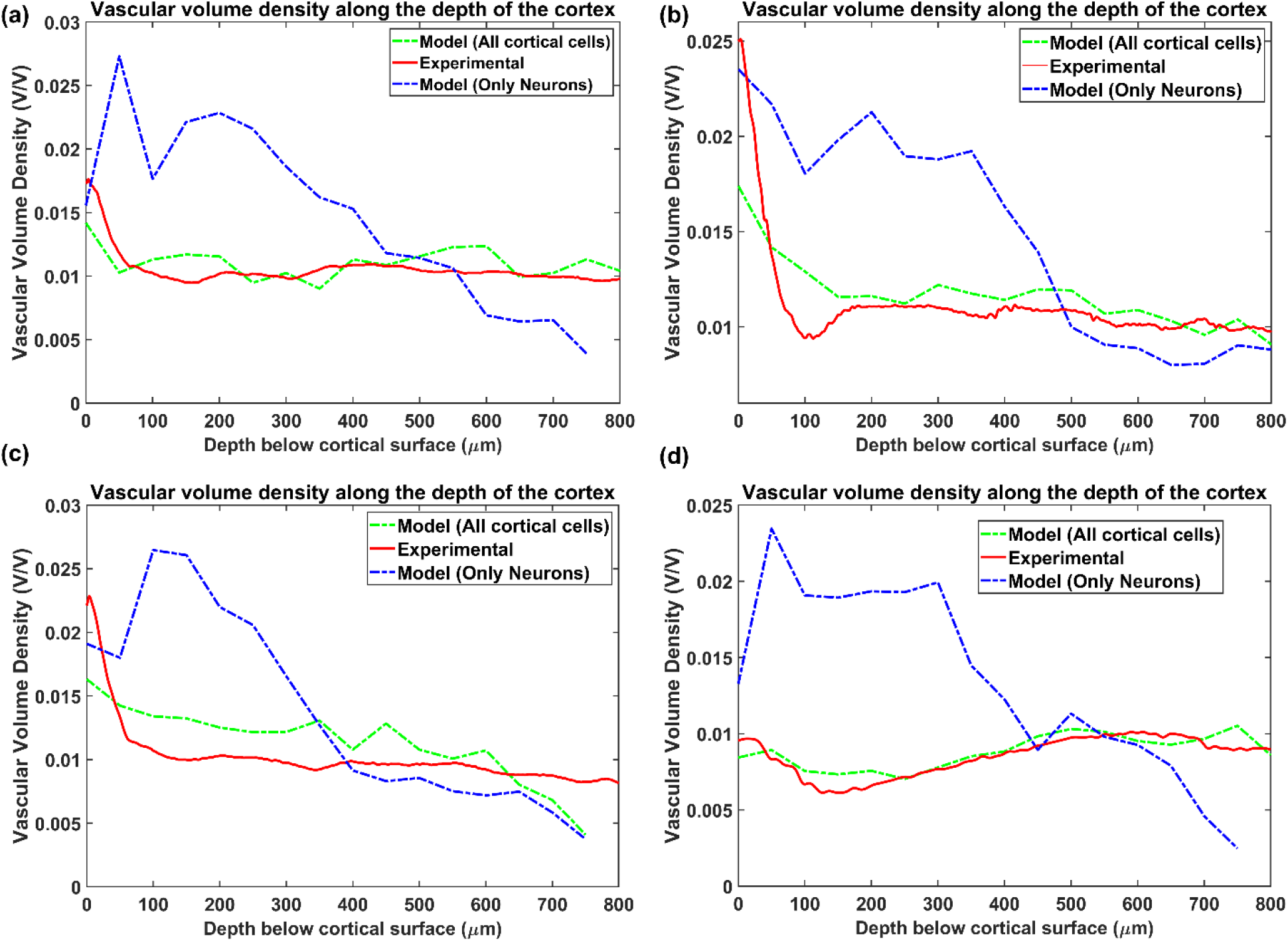
Comparison of the vascular volume density from all the four regions when considering neurons alone and both neurons and non-neural cells. The four regions are (a) Agranular cortex which was measured 1mm anteroposterior and 1mm mediolateral of the Bregma, (b) Partially granular cortex measured −1 mm anteroposterior and 1 mm mediolateral of the Bregma, (c) Partially granular cortex measured 2 mm anteroposterior and 1 mm mediolateral of the Bregma (d) Granular cortex measured 1 mm anteroposterior and 4 mm mediolateral of the Bregma. The experimental data is from Tsai et al 2009 [28].

The figs. 8 a-d show the reconstructed microvascular volume density and the total vascular volume density in comparison with the experimental values. We defined the vessels of radii less than 3μm[28] (all the vessels below level 7) as part of microvasculature. The similarity of the model with the experiment shows that the radii distribution along the levels and vascular architecture is a reasonable approximation of the biologically observed vasculature.

**Figure 8:**
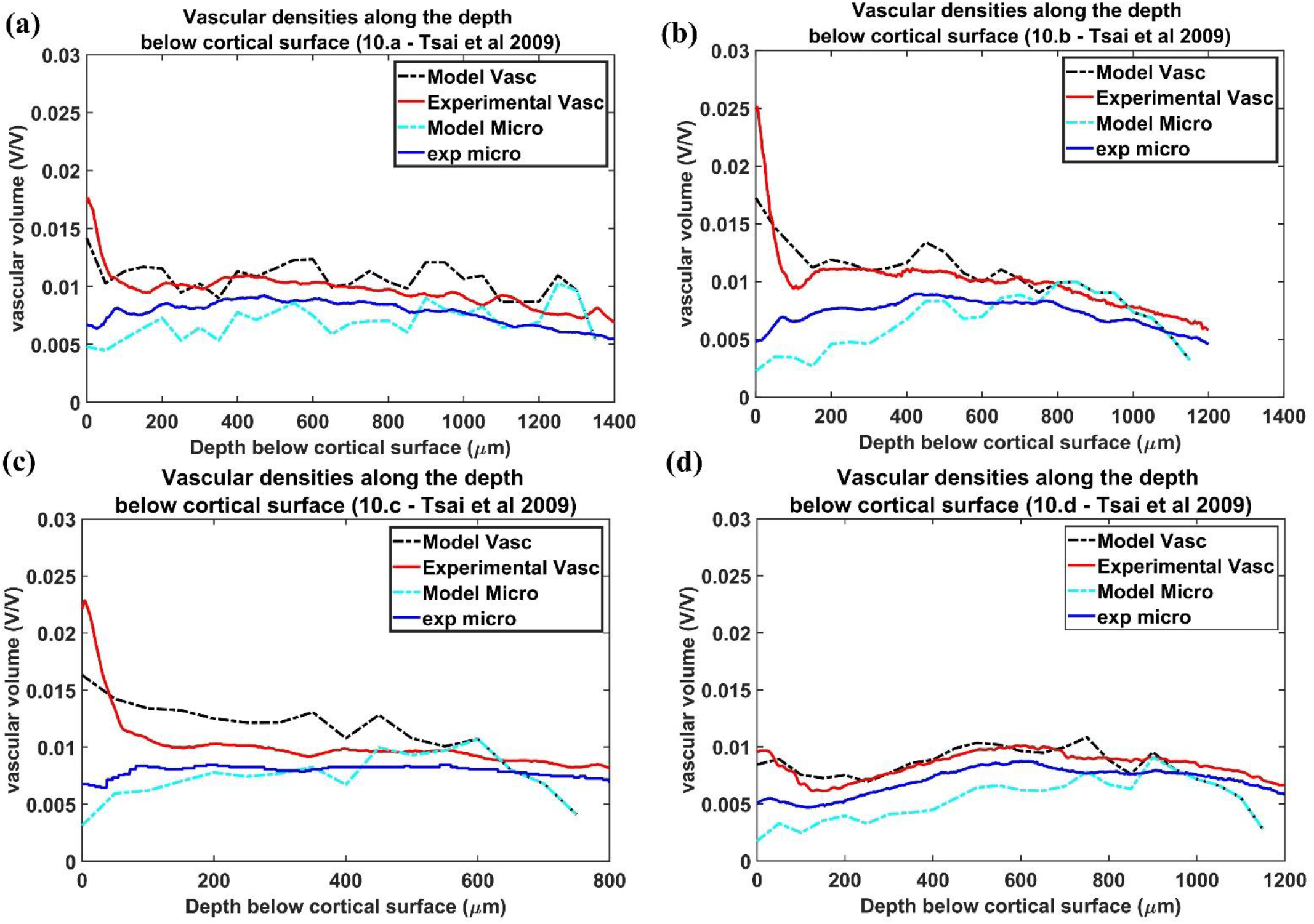
Estimation of vascular volume density and microvascular volume density from all the four regions. The red plot and the blue plot shows the experimentally observed total vasccular volume density and microvascular volume density respectively. The black and the cyan plots trace the values of total vascular volume density and microvascular volume density as simulated by the model. The four regions are, (a) Agranular cortex which was measured 1mm anteroposterior and 1mm mediolateral of the Bregma, (b) Partially granular cortex measured −1 mm anteroposterior and 1 mm mediolateral of the Bregma, (c) Partially granular cortex measured 2 mm anteroposterior and 1 mm mediolateral of the Bregma (d) Granular cortex measured 1 mm anteroposterior and 4 mm mediolateral of the Bregma

### Mean distance between vessels and neurons

The mean distance between vessels and neurons was estimated for each layer as per eqn. (19) and the distribution was depicted as a histogram. Figs. 9(a-d) shows the histogram of mean distance between neurons and the closest vessel calculated across the layer. It was observed that in all the four cases, the mean value is between 0.010 mm and 0.013 mm, which is close to 0.014mm, the biologically observed value as shown in fig 9.c of [28].

**Figure 9:**
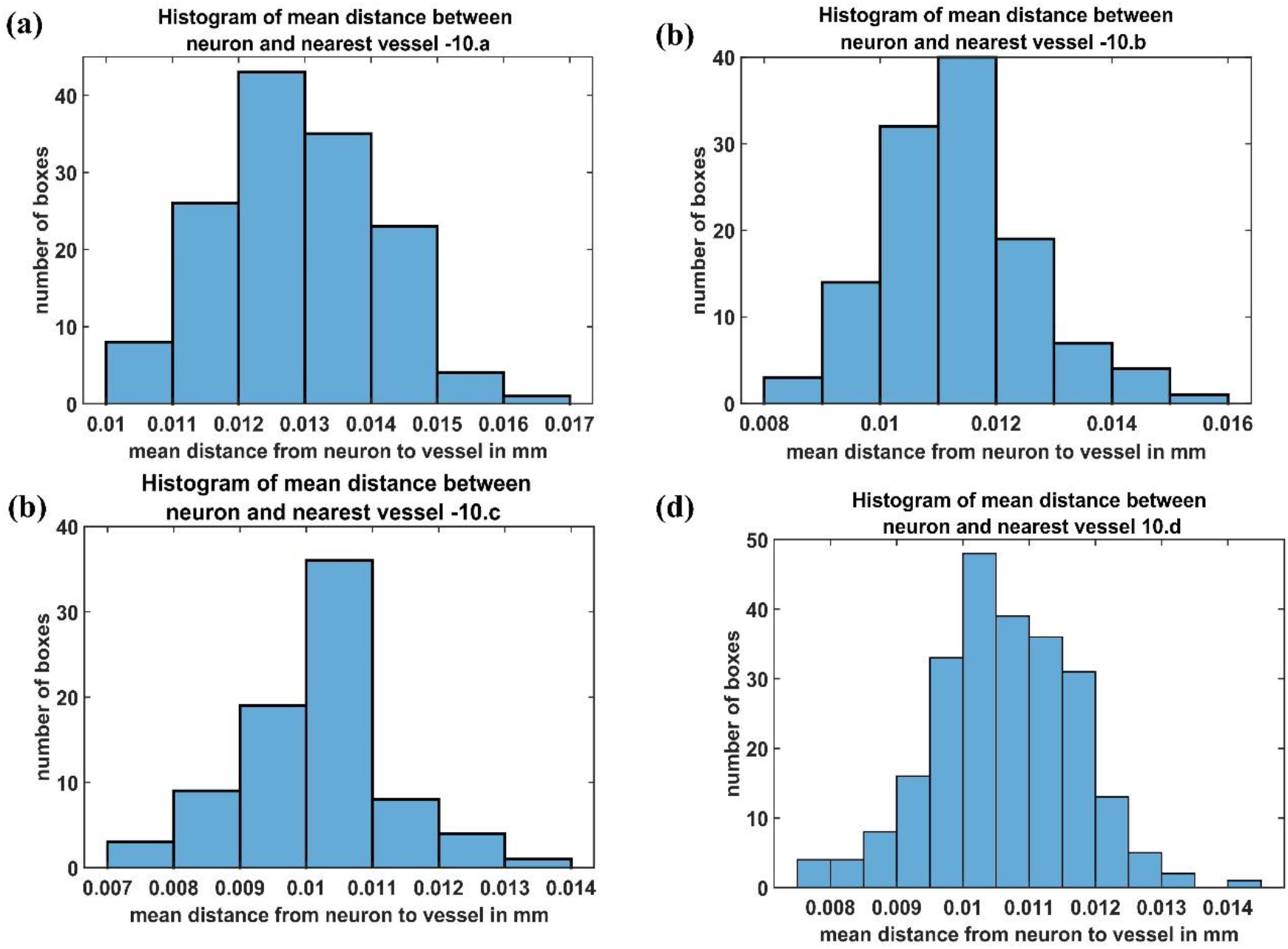
The histogram of mean distance between neurons and the nearest vascular leaf nodes. The measurements are from four regions namely (a) Agranular cortex which was measured 1mm anteroposterior and 1mm mediolateral of the Bregma, (b) Partially granular cortex measured −1 mm anteroposterior and 1 mm mediolateral of the Bregma, (c) Partially granular cortex measured 2 mm anteroposterior and 1 mm mediolateral of the Bregma (d) Granular cortex measured 1 mm anteroposterior and 4 mm mediolateral of the Bregma

### The correlation between neural areal density and vascular volume density

Direct visual inspection of the neural density distribution (fig.4) and vascular density distribution (fig.7 and fig.8) suggests that the neural density does not covary with the vascular density, agreeing with experimental studies. To confirm this inference, the average Pearson correlation coefficient was also estimated and found to be less than 0.2. We then checked if the model was able to show the correlation which exists between neural areal density and microvascular density when compared over a global scale (across species and regions). We simulated the interspecies variation by choosing the number of vascular leaf terminals randomly between 35% to 45% of total neurons for each trial. Along with this, the random distribution of the neural terminals inside each layer also provided randomness. We collected data from 25 simulations in total for the four regions, and used the combined data to analyse the correlation. The neural areal density was calculated as the density of the neurons in the region under consideration multiplied by the depth of the region. As observed in experimental study by Tsai et al (2009 [28], we saw that the model also predicts a correlation of 0.61. The scatter plot between neural areal density and microvascular volume density across different animal models and regions is shown in fig.10.

**Figure 10:**
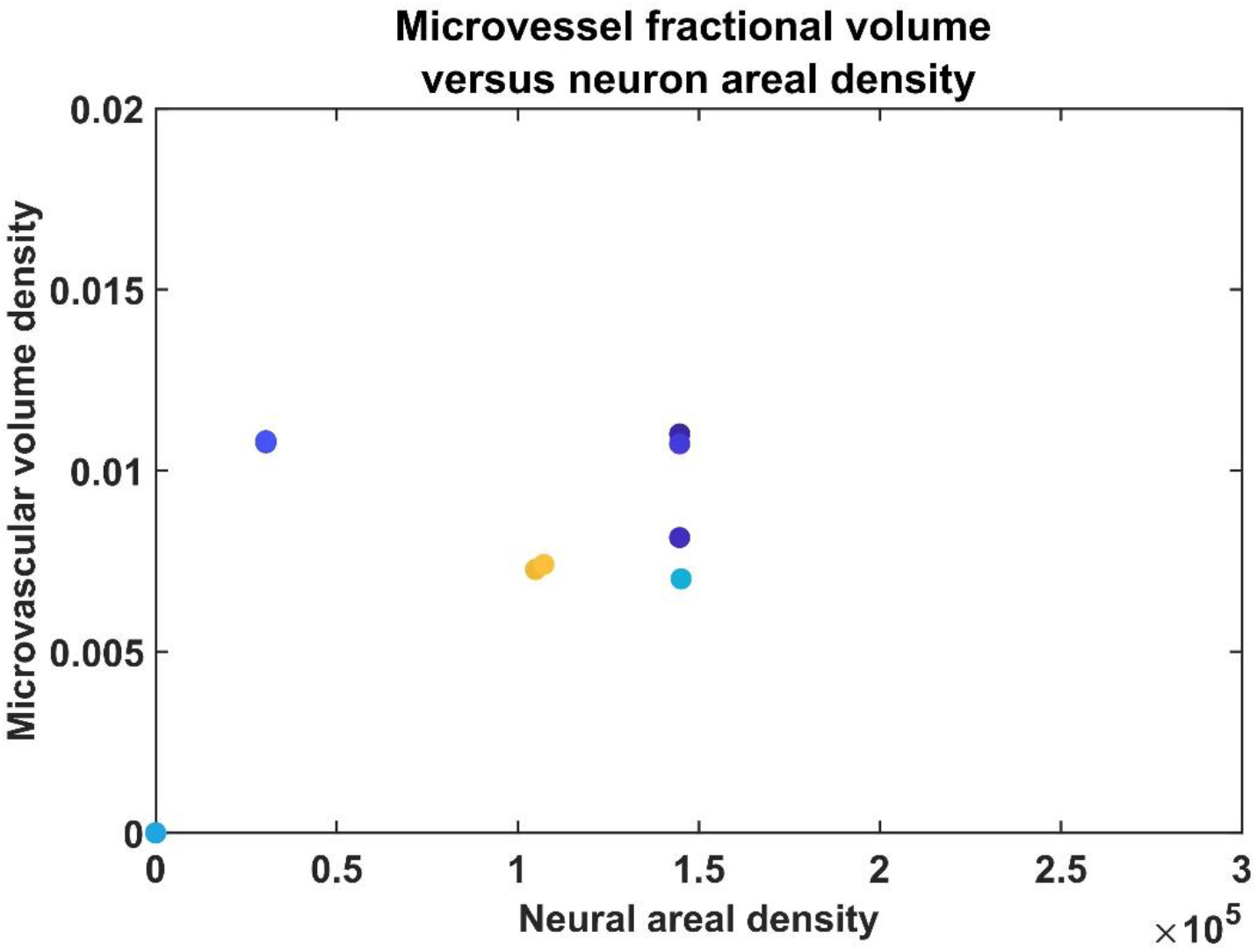
The scatter plot between neural areal density and microvascular volume density of different animals

### Influence of neural activity on vascular arborization

Once the VAM study established the influence of the cytoarchitecture of neural and non-neural cells on vascular arborization, we explored whether neural activity could also influence the arborization of vessels. We modified the VAM to the A-VAM to incorporate the neural activity. In order to consider the influence of neural activity on vascular arborization, a larger area of cortical tissue needs to be considered (spanning multiple barrels in the whisker barrel cortex). The vascular arborization in a larger area (0.6 mm X 0.6 mm X 0.7 mm) was hence considered to model A-VAM. The neural and non-neural cells were distributed similar to the VAM implementation as shown in fig. 11.

**Figure 11:**
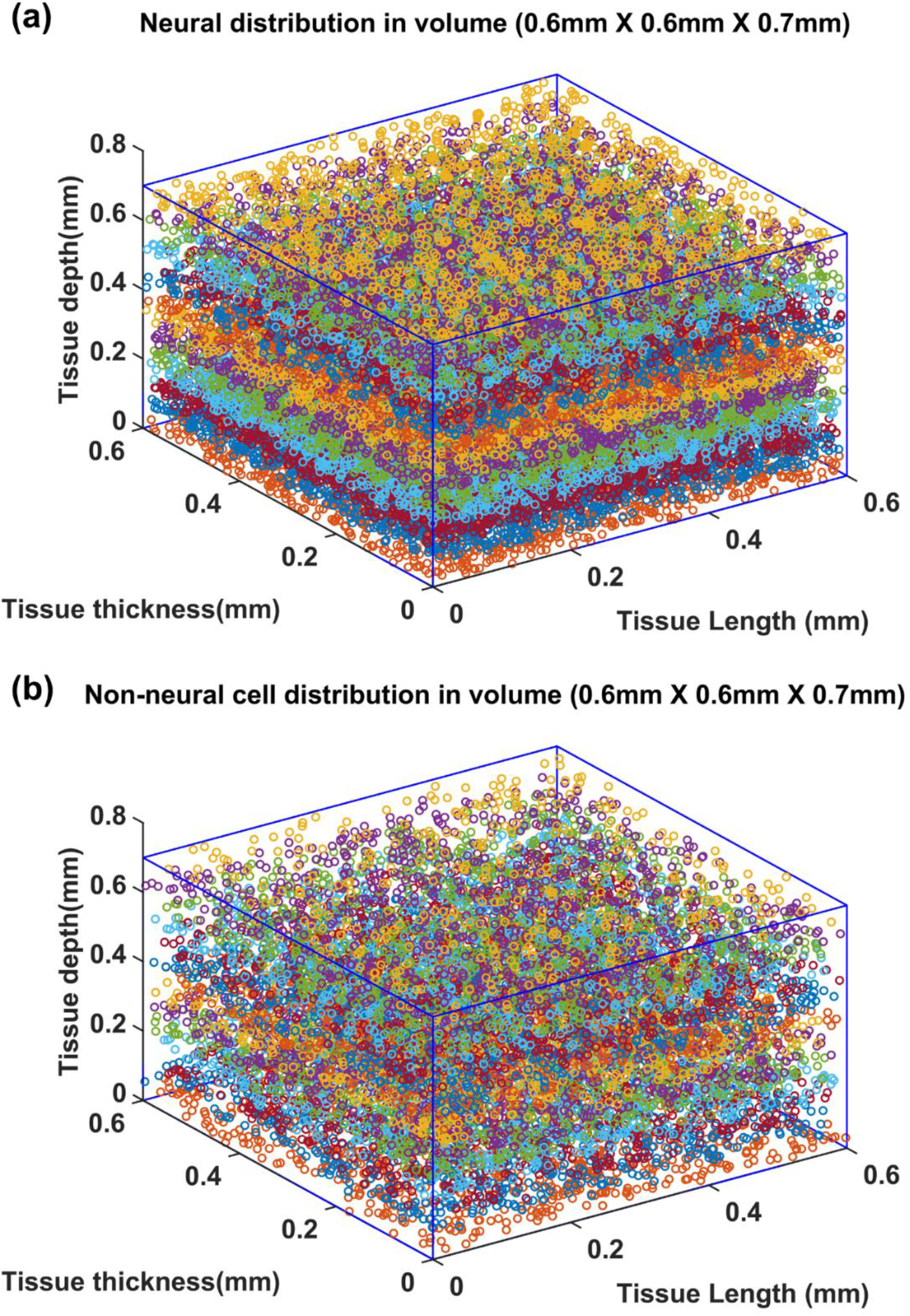
Distribution of (a) neural and (b) non neural cells in a larger volume (0.6mmX0.6mmX0.7mm).

**Figure 12:**
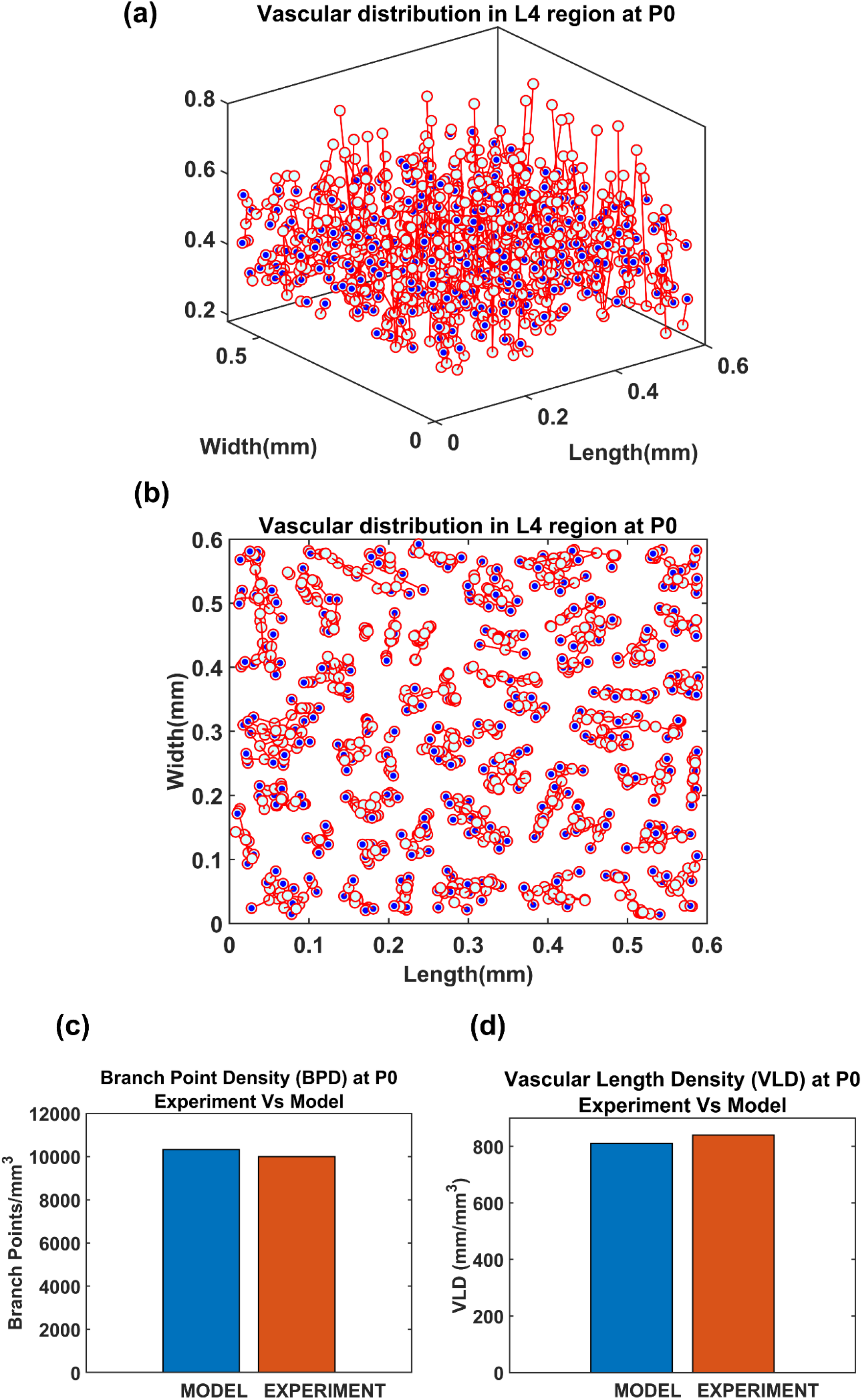
Vascular distribution at the L4 region at the time of birth (P0). The blue dots denote the vascular terminal leaf nodes. (a) The 3D view of the L4 region (b) The top view of the L4 region.

Since a larger area was considered in this case, the vascular growth was initialized by defining 44 root nodes [29] distributed over the surface of the cortical tissue volume. The vascular arborization begins in the prenatal growth phase where the vascular growth depends only on the cytoarchitecture of the tissues until birth. The arborization at birth in a small volume (L4 region, between 0.3mm and 0.55mm [29]) as shown in fig. 11.

At P0, the neural activity is incorporated by means of connecting a SOM network to the VAM (fig. 3) making it the A-VAM. The SOM was trained for the entire whisker barrel cortex as shown in fig. 13(a). Each coloured blob represents one barrel and its activity corresponds to the activity of the whisker which it represents. The SOM interpixel distance was defined true to the biological whisker barrel dimensions [6]. An area of 0.6mm X 0.6mm was chosen (fig 13.b) which contained three barrels each of B, C and D row whiskers (Enclosed by red square in fig 13.b) for incorporating with the VAM.

**Figure 13:**
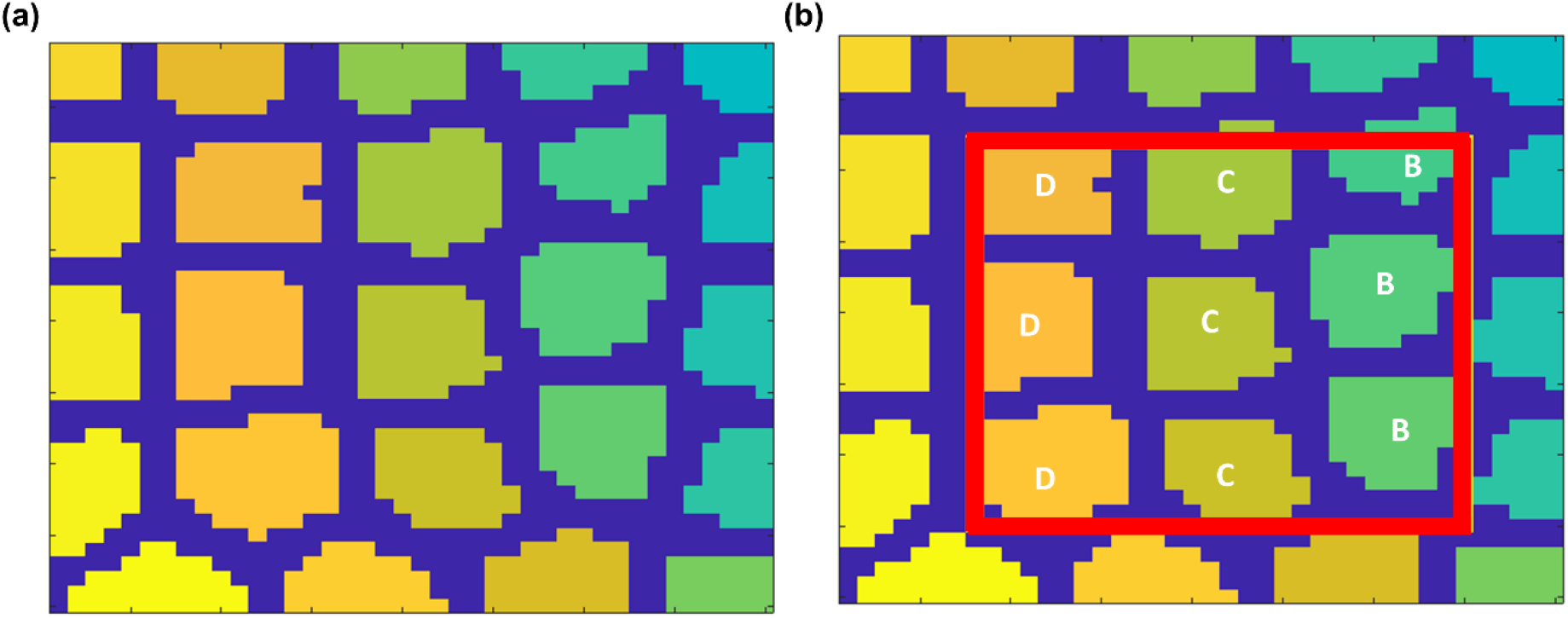
SOM trained to represent whisker barrel cortex: (a) Full whisker barrel cortex. (b) The 0.6mm X 0.6mm area corresponding to the cortical tissue considered for A-VAM

The selected area of SOM network was stacked on the L4 area. The neural activity of each cortical neuron was estimated based on the activity of the nearest SOM neuron, and based on how far the cortical neuron is from that nearest SOM neuron. The weights of SOM during various stages of training were saved and the output of the SOM was systematically presented to the A-VAM network at every 25^th^ iteration. We adopted such a method to ensure simultaneous learning in neural and vascular network without much computational load. Since vascular feedback to neurons is not incorporated in the current model, such an approach can be justified.

The neural activity dependent vascular arborization was explored under two criteria: (a) Control – where all the nine whiskers in the selected region were intact; (b) Lesioned - where the middle row (C row) whiskers were lesioned. In the control model, all the whiskers are represented by approximately the same area in the SOM as shown in fig. 14a. In the lesioned model, the representation of whiskers in the trained SOM network look like in fig. 14b. The area representing the lesioned middle row (C row) shrinks while the neighbouring rows (B and D) are represented by a larger area. With the help of A-VAM, we explore how such a change would affect the vascular arborization. The yellow lines in both fig. 14a and fig. 14b mark the boundary of the C row considered to evaluate the vascular density.

**Figure 14:**
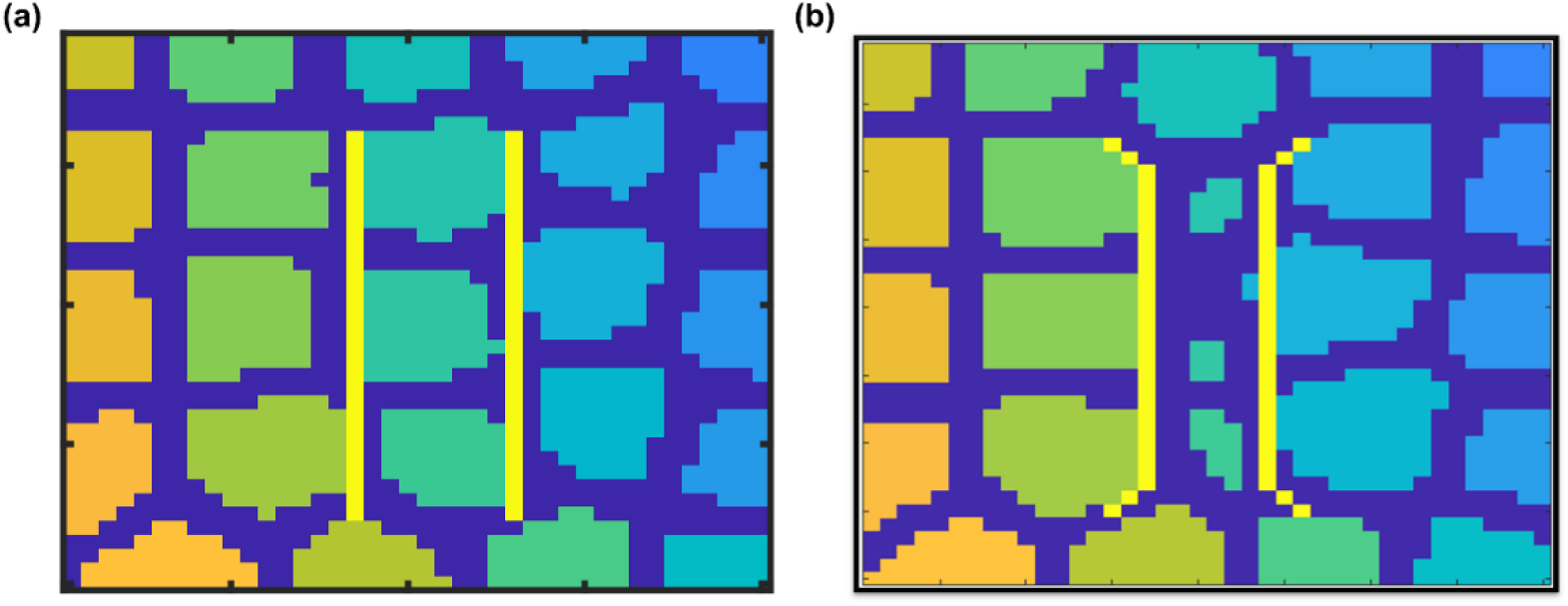
Boundary of C row barrels in (a) Control model (b) Lesioned Model

The vasculature was allowed to grow, and span the tissue region until the tree converges (until all growing vascular nodes are declared as terminal leaf nodes). The settled vascular tree in the L4 region of the control and lesioned cases were as observed in figs. 15a and 15b. The top views of the 3D plots are shown in figs. 15c and 15d. The blue dots represent the vascular leaf nodes. By visual inspection of the top views of the L4 region (figs. 15c,d), we can see that there is a reduction in branch point density in the lesioned model when compared with the control model. To get a more quantitative account of the difference, we calculated the BPD and VLD of both the models at the L4 volume in the area between the yellow lines marked in figs. 14a,b. The resulting values showed that the BPD and VLD of the lesioned model was significantly lesser when compared to the control model, as observed by the experimental study. This concludes that the neural activity along with the cytoarchitecture of neural and non-neural cells influences vascular arborization.

**Figure 15:**
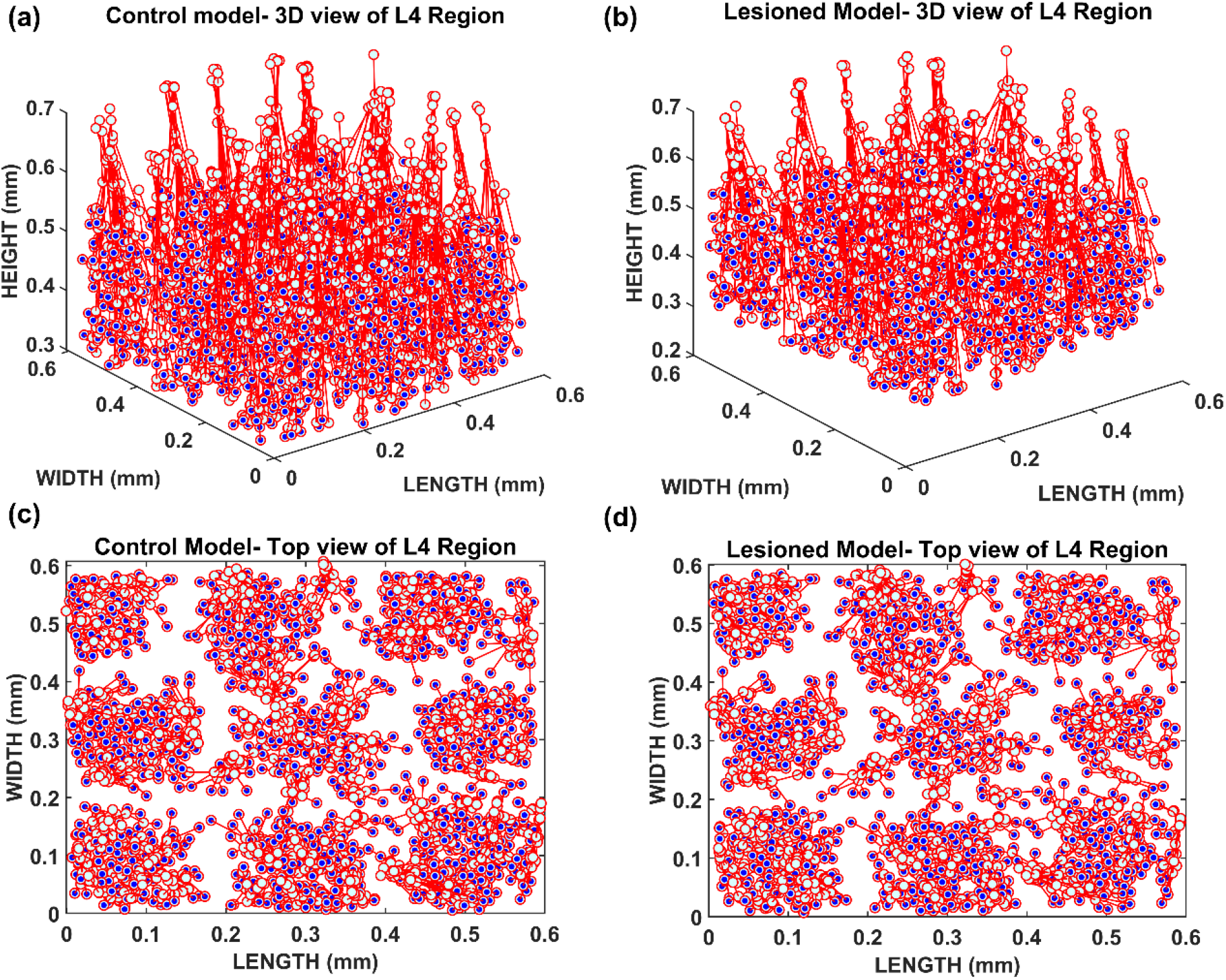
The vascular arborization in L4 region of whisker barrel cortex

**Fig 16:**
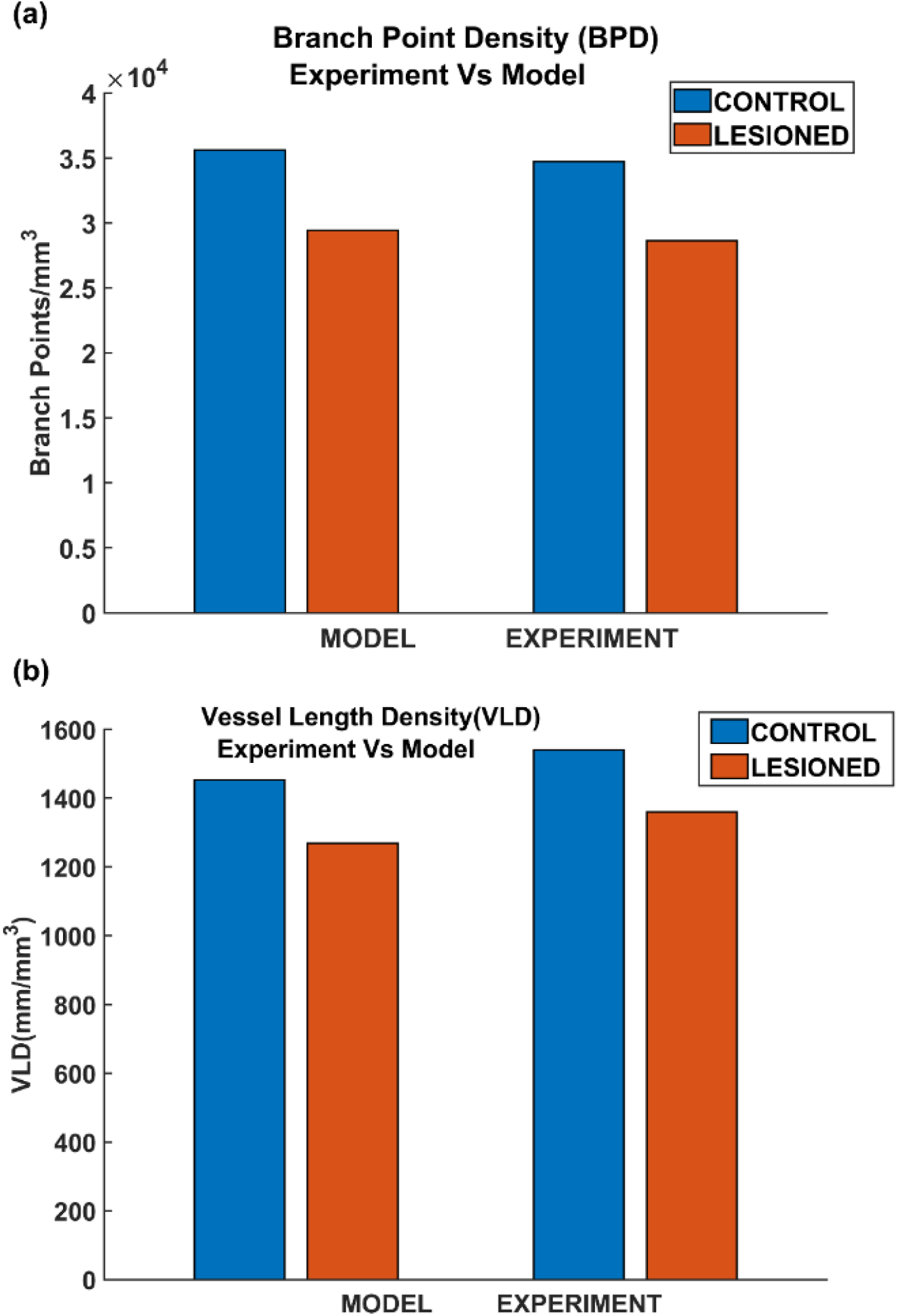
Comparison of branch point density (BPD) in control and lesioned animal. (b) Comparison of vessel volume length density (VLD) in control and lesioned model

## Discussion

Our first model of vascular tree growth, VAM, which was based purely on the cytocarchitecture of the cortical cells, was able to simulate a fully grown vascular network from a single root node. The model resulted in a vascular tree that closely approximates the experimental vascular tree as characterized by several indices. The microvascular volume density and the total vascular volume density estimated from the model were in close agreement with the experimental studies [28]. The mean distance between vessels and neurons in the model was close to 14 μm, the biologically observed value. The model also shows that the neural densities do not covary with the vascular density along the depth of the cortex, but revealed a high correlation between neural areal density and microvascular density when compared over a global scale (across animals and regions). Moreover, the range of estimated vessel radii that resulted in this vascular volume density agreed with the range of arteriolar diameters observed experimentally in rats and mice [30,31].

Inspired by the recent observations that the neural activity also could influence the vascular arborization, we modified the VAM to A-VAM, making the tree growth depend on the neural activity as well. The neural layer modelled using SOM network impacted the vascular tree growth in both the growth phase and settling phase. The grown tree was characterized in terms of branch point density and vascular volume density. The vascular arborization observed in control vs. lesioned models evidently showed that the branch point density and the vascular volume density are significantly lower in the latter. This was in agreement with the experiments done on mice whisker barrel cortex [6].

The study corroborates the statement that neurovascular coupling begins from a very early stage of growth. The development of neural and vascular networks takes place in parallel such that there is a mutual influence. The comprehensive understanding of the influence of neural activity on vascular arborization could be a game-changer in stroke rehabilitation therapy. Angiogenesis and neurogenesis post-stroke conditions are known to be mediated by a number of molecules of both neural and vascular origin [32–36]. Even though the relationship between angiogenesis and neurogenesis in the ischemic brain is theoretically hypothesized, it is yet to be validated. In such a scenario, a bidirectionally coupled neurovascular growth model would help to understand the significance of multiple pathways which are hypothesized to play a role in neurovascular remodelling.

The role of computation in shaping the morphological characteristics of neuronal branching was investigated by Cuntz et al [19] based on the computational principles that govern neuronal morphologies laid out by Ramon Y Cajal [37]. They showed that a simple growth algorithm that optimizes the total cable length and the path length from any point to the root in an iterative fashion can generate synthetic dendritic trees that are indistinguishable from their real counterparts for a wide variety of neurons.

The astrocytic arborization is relatively less explored. A recent study [38] modelled the astrocyte morphology by forming Voronoi partitioning around randomly distributed seed points and matching the polygonal area patch from a given Voronoi partition to the closest astrocyte template extracted from experimental images of astrocytes. However, to the best of our knowledge, there are no published computational models that simulate the arborization of astrocyte processes.

Neurons are dependent on the continuous supply of energy substrates from the cerebral vessels, and hence their firing depends on the vascular feedback to a large extent. However, in case of the A-VAM, though it captures the neural influence on angiogenesis, the current version does not include the vascular influence on neural activity. For the initial validation, we assume a healthy vascular supply and hence the feedback to the neurons were assumed to be adequate and continuous.

The relevance of a healthy vasculature in shaping the function of the neural network has been discussed extensively [39–46]. Neurodegenerative diseases are being traced back to an impaired neurovascular functional coupling [1,47–50]. As opposed to the conventional view of neurons as the primary computational unit of the brain, we postulate that the neurovascular unit as a single combined unit is a more appropriate computational unit of the brain. We conclude by emphasizing that the coupling between neural and vascular networks is not limited to the functional aspects but also decides the structural development of both networks. To conclude with a poetic comment by Bautch and James [7] – neurovascular coupling is indeed a beautiful friendship that begins in the womb and continues till the end.

## Acknowledgments

We thank The Department of Biotechnology (DBT), Ministry of Science and Technology, Government of India (BIO/17-18/303/DBTX/SRIN), and the Center for Complex Systems and Dynamics, IIT Madras for partial funding of this project.

## Author Contributions

BSK did model designing, coding, analysis of the results, and manuscript preparation, SM did model tuning, analysis of results and manuscript preparation. SRG did model tuning and analysis of results. VSC did model designing, analysis of the results, and manuscript preparation. All authors commented on and approved the final manuscript.

