## Supplementary Information for "The Influence of Neural Activity and Neural Cytoarchitecture on Cerebrovascular Arborization: A Computational Model"

#### S1: Calculation of length of vessel in each layer

The length of each vessel (L), location of its parent node (P1) and child node (P2) and the depth of each layer is known. An example is considered in fig. S1. Let the total length of the vessel be given by

$$L_T = L1 + L2 + L3 \quad \text{(S1)}$$

From the knowledge of depth of each layer and the coordinate of beginning and end of the vessels, the vertical distances can be calculated between (i) P1 and the end of the first layer and is denoted by Z1, (ii) Depth of the second layer (Z2) (iii) distance between end of the second layer and P2 denoted by Z3.

Applying trigonometric principles

$$L1{:}\,L2{:}\,L3 = Z1{:}\,Z2{:}\,Z3 \quad \text{(S2)}$$

Assuming K to be a constant,

$$(Z1 + Z2 + Z3)K = L_T \quad \text{(S3)}$$

$$K = \frac{L_T}{(Z1 + Z2 + Z3)} \quad \text{(S4)}$$

$$L1 = KZ1 \quad \text{(S5)}$$

$$L2 = KZ2 \quad \text{(S6)}$$

$$L1 = KZ3 \quad \text{(S7)}$$

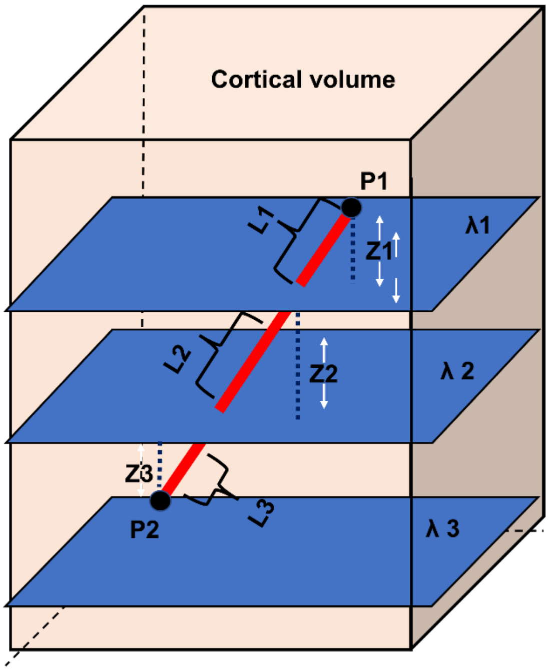


Figure S1: Calculation of length of vessel segment in each layer

### S2: Self-Organizing Maps (SOM)

Self-Organizing Maps (SOM) was introduced by Teuvo Kohonen in the 1980s. The fundamental idea behind the SOM is the feature learning facilitated by competitive learning among a network of neurons.

1. Initialize SOM with random weights or using the data itself
2. Give an input and search for the maximum response neuron
3. Select the neighborhood size and a learning rate (η)
4. Update the weight ($W_p$) of the $p^{th}$ neuron such that

$$\Delta w_p = \eta \Lambda (X - W_p) \tag{S8}$$

$$\Lambda(p, q) = e^{--\frac{\|p-q\|^2}{\sigma^2}} \tag{S9}$$

   Where p is the index of winning neuron, q is the index of the neighbor neuron, $\sigma$ is the standard deviation of the Gaussian function.

5. Repeat 4 till the weight converge.

   **Parameters used in the model to train SOM:**

   η=0.1

   $\sigma = 4$

   neighborhood radius=20
